# Adaptation of utility functions to reward distribution in rhesus monkeys

**DOI:** 10.1101/2020.05.22.110213

**Authors:** Philipe M. Bujold, Simone Ferrari-Toniolo, Wolfram Schultz

**Affiliations:** Department of Physiology, Development and Neuroscience, University of Cambridge, Cambridge CB2 3DY, United Kingdom

## Abstract

This study investigated the influence of experienced reward distributions on the shape of utility functions inferred from economic choice. Utility is the hypothetical variable that appears to be maximized by the choice. Despite the generally accepted notion that utility functions are not insensitive to external references, the exact occurrence of such changes remains largely unknown. Here we benefitted from the capacity to perform thorough and extensive experimental tests of one of our evolutionary closest, experimentally viable and intuitively understandable species, the rhesus macaque monkey. Data from thousands of binary choices demonstrated that the animals’ preferences changed dependent on the statistics of recently experienced rewards and adapted to future expected rewards. The elicited utility functions shifted and extended their shape with several months of changes in the mean and range of reward distributions. However, the adaptations were usually not complete, suggesting that past experiences remained present when anticipating future rewards. Through modelling, we found that reinforcement learning provided a strong basis for explaining these adaptations. Thus, rather than having stable and fixed preferences assumed by normative economic models, rhesus macaques flexibly shaped their preferences to optimize decision-making according to the statistics of the environment.

## Introduction

Every day we make choices between outcomes that vary widely, sometimes on the order of magnitudes. In a single morning, we can go from choosing between outfits, to choosing to visit our favourite cafe, to comparing the costs of a train or plane journey for our next holiday destination. Yet, despite the complexity of representing all of these situations, we manage - with a relatively limited brain - to mentalise and indeed optimise the majority of our choices.

Prospect Theory (PT), the dominant model in behavioural economics, posits that we optimize our decisions by calculating the value of our choices relative to a reference-point (Kahneman & Tversky, 1979; Tversky & Kahneman, 1986). That is, rather than objectively evaluating the outcome of our choices, we perceive our options as gains or losses depending on what we are expecting: if an outcome is better than our reference, we treat it as a gain; if is is worse, we treat it as a loss. Mathematically, PT represents this behaviour with an S-shaped value (or utility) function where the subjective value of gains and losses is given by concave and convex parts of the function, respectively. This has important behavioural consequences, particularly for risky-decision-making, as this normative (utility) framework predicts that people’s tendency to make risk averse decisions depends on their perception of outcomes as being gains or losses.

While the idea of reference-dependence has been readily adopted by modern decision theory (Rabin, 2000; Wakker, 2010), economists are still unclear about how reference points form (Barberis, 2012). In prospect theory (PT), Kahneman and Tversky abstractly define reference-points as exogenous from the decisions being made. That is, the reference point is not directly explained by PT and can be shaped by “*aspirations, expectations, norms, and social comparisons*” (A. Tversky & Kahneman, 1991, p.157). Alternatively, recent economic models consider reference points an epiphenomenon of the way in which our mind adapts to the statistics of the task at hand (Delquié & Cillo, 2006; Köszegi & Rabin, 2006; Sugden, 2003) - a framework more in line with the findings that, far from being restricted to human reasoning, reference-dependence is a homogeneous feature of primate decision-making (Santos & Rosati, 2015) and the brain (Carandini & Heeger, 2012; Louie et al., 2013; Padoa-Schioppa, 2009; Tremblay & Schultz, 1999). Along these lines, one particularly interesting proposal from the epiphenomenon framework is that of range-dependent utility, or RDU (a play on reference-dependent utility; see Kontek & Lewandowski, 2018). Inspired by psychology’s *range-frequency theory* (Parducci, 1965, 2012) and neurobiology’s *efficient-coding hypothesis* (Laughlin, 1981; Summerfield & Tsetsos, 2015), RDU suggests that decision-makers evaluate the value of their options relative to not one, but two reference points: the minimum and maximum rewards available in any given scenario. In this view, what PT identifies as a referencepoint could be nothing more than the product of a utility function that adapts to the distribution of possible rewards: the point at a sigmoidal curve inflects from convex to concave (mimicking a neuron’s tuning curve; Carandini & Heeger, 2012; Webster, Werner, & Field, 2005).

Because studies on reference-dependence generally focus on identifying a unique reference-point (Baillon et al., 2015), or on describing behaviours under specific reference predictions (Allen et al., 2016; Crawford & Meng, 2011; Wenner, 2015), there is, as of yet, no way of corroborating or contradicting the previous hypotheses on the emergence of reference-points. The few studies that consider shifts in preferences generally do so in a single distribution, local context: they document reference-point changes following the wins or losses of risky gambles (Arkes et al., 2008, 2010; Shi et al., 2015); never the impact that changes in expectation have on decision-making. Concurrently, little is known about the impact of a task’s structure on preferences, nor how different reward statistics might translate to reference-points.

Animal experiments allow far higher trial numbers and longer experimental timescales than human studies do, they allow us to explore the formation of reference-points both in utmost detail and in subjects where exogenous factors have minimal impact (i.e. no contribution of language or higher numerical ability). To that effect, we investigated how the reward distribution experienced in a binary choice task - defined on different reward magnitudes and spreads - shaped the preferences of rhesus macaques (a species that displays many, if not most, of the fundamental choice patterns humans display; Heilbronner & Hayden, 2013, 2016; Stauffer et al., 2015). we presented macaques with several sets of risky choice options in which the distribution of reward magnitudes remained stable for weeks at a time, then suddenly shifted to a new distribution (higher/lower magnitudes or wider/narrower spread). On each testing day, we fit the animals’ choices with S-shaped utility functions that could explain both risk seeking and risk averse choices (Genest et al., 2016; Stauffer et al., 2014). We then looked at how the animal’s risk preferences changed as a function of the reward distribution they experienced. We found that, while utilities stayed relatively put for periods during which a single reward distribution was experienced, the animals consistently shifted their preferences when a novel reward distribution was introduced. In fact, the shape of estimated utility functions mirrored the lowest and highest rewards that monkeys had experienced over the course of the preceding weeks – even if these now fell outside of possible. From these findings, we suggest that far from being fixed and abstract, preferences follow the expectation of what animals think might happen given the knowledge they have accumulated over time.

## Methods

### Animals

This research has been ethically reviewed, approved, regulated and supervised by the following UK and University of Cambridge (UCam) institutions and individuals UK Home Office, implementing the Animals (Scientific Procedures) Act 1986, Amendment Regulations 2012, and represented by the local UK Home Office Inspector, the UK Animals in Science Committee, the UK National Centre for Replacement, Refinement and Reduction of Animal Experiments (NC3Rs), the UCam Animal Welfare and Ethical Review Body (AWERB), the UCam Biomedical Service (UBS) Certificate Holder, the UCam Welfare Officer, the UCam Governance and Strategy Committee, the UCam Named Veterinary Surgeon (NVS), and the UCam Named Animal Care and Welfare Officer (NACWO).

Three male rhesus macaques (Macaca mulatta) weighing 11.2, 15.3, and 13.2 kg (Monkeys A, B and C, respectively) participated in this experiment. All animals used in the study were born in captivity, at the Medical Research Council’s Centre for Macaques (CFM) in the UK. The animals were pair-housed for most of the experiment; monkeys B and C shared an enclosure. The animals ranged in age from 5 to 8 years old, and all subjects had previous experience with the visual stimuli and experimental setup (Ferrari-Toniolo et al., 2019).

### Behavioural task and training

Rhesus monkeys are the most commonplace species of non-human primate found in scientific research (Capitanio & Emborg, 2008). There is therefore a rich literature reproducing human economic choices in rhesus macaques. Most relevant here is that rhesus macaque behaviour can be successfully predicted using PT (Farashahi et al., 2018; Ferrari-Toniolo et al., 2019; Genest et al., 2016; Stauffer et al., 2015). In addition, macaque experiments allow us to control the pre- and post-experimental environments in ways not possible for human studies – we can ensure that experimental variables are independent of rewards and choices made outside of the experiment (Chen et al., 2006). For this study, the delivery and distribution of rewards experienced were unique to the experimental setup. The animals experienced nothing comparable outside of the laboratory.

Each animal used a left-right joystick (Biotronix Workshop, University of Cambridge) to make choices between reward-predicting stimuli presented on a computer screen. After each choice, the animals received their chosen reward in the form of a specific blackcurrant juice quantity delivered probabilistically (matching the probabilities indicated by each stimulus).

The animals were presented with a simple visual stimulus consisting of one or two horizontal lines positioned inside a frame of two vertical lines depicting reward options that varied both in magnitude (i. e. liquid quantities, ml) and in the probability of a reward being delivered. Reward magnitudes were represented by the vertical position of the horizontal lines on the screen, whereas reward probability was represented by the lenght of the horizontal lines inside the framing lines (Fig. 1a). Safe (riskless) options were represented by singular full-width horizontal lines that touched both sides of the frames, whereas gambles with multiple risky rewards were signalled by multiple horizontal lines within the vertical frame.

**Figure 1.**
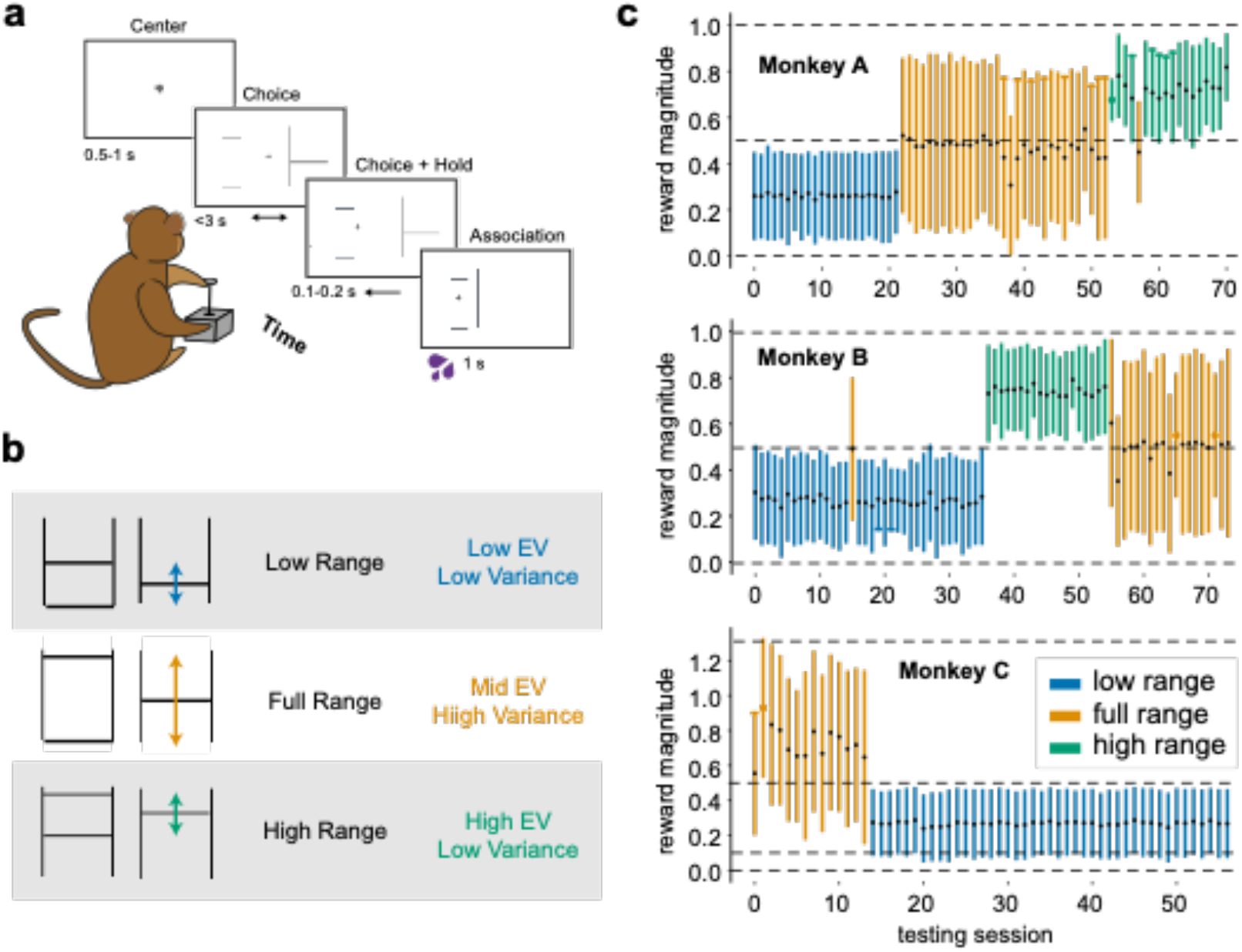
Experimental design and timescale. a) Binary choice task. The animals chose one of two gambles with a left-right motion joystick. They received the blackcurrant juice reward associated with the chosen stimuli after each trial: the reward’s magnitude and probability of delivery were signalled by the vertical position and width of a horizontal line as set between two vertical ones. Times, in seconds, indicate the duration of each of the task’s main events. b) Experimental reward distributions. Choices were made in one of three experimental reward distributions. In the low distribution, choice options had juice magnitudes set between 0 ml and 0.5 ml during preference elicitation sequences. The high distribution involved juice magnitudes set between 0.5 ml and 1.0 ml during preference elicitation sequences (unique to Monkey A and B). The full distribution was set between 0 ml and 1.0 ml for Monkeys A and B and set between 0.1 ml and 1.3 ml for Monkey C. c) Monkeys’ experienced specific reward distributions for consecutive days. Vertical lines represent the daily experimental session, in their tested order; the height of these lines signals the reward distribution tested (blue, low distribution; yellow, full distribution; green high distribution). Black dots indicate the mean magnitude of all rewards experienced on the day, the white dots represent the standard deviation on the mean.

The animals were trained to associate these two-dimensional visual stimuli with blackcurrant juice rewards over the course of >10,000 single-outcome, imperative trials. In these trials, a single reward option was presented on either the left or right side of the screen. To obtain the cued reward, the animals were required to select the side on which the reward was presented. After imperative training, where only one option was presented, all experimental data were gathered within a binary choice paradigm in which the animals chose one of two reward options presented simultaneously. One option was always a gamble; the other was always a safe, guaranteed reward. Every choice trial began with a white cross at the centre of a black screen, followed by the appearance of a joystick cursor. To initiate a trial the animal had to move the joystick cursor to the center cross and hold it there for 0.5-1s. After a successful central hold, two reward options appeared to the left and right of the central cross (Fig. 1a). The animal had 3s to convey its decision by moving the joystick to the side of its choice and holding it there for 0.1s to 0.2s, after which time the unselected option would disappear. The selected option remained on the screen for 1s after reward delivery to strengthen any stimulus-reward associations with visual feedback. A variable intertrial period of 1–2 s (blank screen) preceded the next trial. Errors were defined as trials with an unsuccessful central hold, trials in which the animal failed to hold the selected side, or trials in which the animal made no choices, and resulted in a timeout of 6 seconds, after which time the trial was repeated.

Reward options were presented in pseudorandom alternation on the left and right sides of the computer screen to control for any side preference. Event times were sampled at 2 khz and stored at 1 khz on a Windows 7 computer running custom MATLAB software (The mathworks, 2015a; Psychtoolbox version 3.0.11), and all further analyses were done using custom Python code (Python 3.7.3, Scipy 1.2.1, Oliphant, 2007). Over the course of 63, 43 and 57 sessions an average of 259 ± 154 (mean ± STD) trials, 317 ± 118 trials, and 131 ± 75 trials were collected for Monkeys A, B and C, respectively. Crucially, animals received the reward they selected after each trial. This ensured that they experienced the rewards they selected with minimal and constant delay, and contrasts with human studies where only a randomly selected subset of trials are rewarded at the end of experimental sessions. Delivering rewards after every trial also allowed us to capture preferences that were contingent on experiences unique to the task - similar delivery method and reward distributions were not experienced in the housing environment.

### Measuring preferences for specific reward distributions

To examine the degree at which preferences are shaped by available rewards, binary choice data were collected from choices between reward options affixed to different reward distributions (Fig. 1b). Three reward distributions were defined in terms of their mean reward magnitude and the spread of possible options i) low-narrow distribution, where tested magnitudes were generally set between 0 ml and 0.5 ml; ii) high-narrow distribution, with magnitudes between 0.5 ml and 1.0 ml; and iii) full distribution, with magnitudes between 0 ml and 1.0 ml (0.1 to 1.3 ml for Monkey C). Importantly, every reward outcome (no matter which distribution) was repeated the same number of times for each session – thus, every reward was equiprobable (flat distribution). We set distributions and kept them fixed for multiple weeks, measuring the effects of reward distribution over weeks rather than blocks in a single session (Fig. 1c). Monkey A experienced a low distribution for 22 days (0 ml to 0.5 ml), a full distribution of rewards for 31 days (0 ml to 1.0 ml), and a high distribution of rewards for 17 days (0.5 ml to 1.0 ml). Monkey B experienced the low distribution for 33 days, then 19 days of high distribution, followed by 18 days of full distribution. Monkey C, quite uniquely, offered a dataset with a longer timescale. He experienced the full distribution of 0.1 ml to 1.3 ml of reward for 14 days then switched to a low distribution of 0 ml to 0.5 ml for 54 weeks. After this, his preferences were measured over 43 days.

Utility functions were estimated for each probability distribution by presenting individual animals with a series of choices between a safe reward (probability of reward, *p* (reward) = 1.0) and a binary, equiprobable gamble (each reward *p* = 0.5) from which Von Neumann–Morgenstern type utilities were estimated. Probability distortions are symmetric and usually minor at *p* = 0.5 (Stauffer et al. 2015; Ferrari-Toniolo et al. 2019); therefore, to obtain utility functions with least fitting errors, we neglected probability distortions and thus assessed EU = *p • u*(ml). To estimate utility functions, we used the fractile-bisection procedure (Machina, 1987), which involves successively dividing the distribution of possible utilities into progressively smaller halves (or fractals) and estimating at each step the magnitude of safe reward at which choices were indifferent against the specific gamble being tested, as done in our laboratory before (Genest et al., 2016; Stauffer et al., 2014). This magnitude is termed certainty equivalent (CE), and represents the subjective value of safe reward that is equivalent to the value of the gamble.

The first step of the procedure involved presenting the animals with choices between this gamble and varying safe rewards (in 0.05 ml increments); in these choices, the safe reward that was equivalent to the gamble in utility terms was identified (i.e. the safe reward chosen in equal proportion to the gamble; see Fig. 2a, b). To estimate this safe reward, the following logistic sigmoid curve was fitted to the proportion of safe choices versus gambles for each of the gamble/safe pairing:

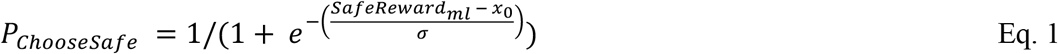

**Figure 2.**
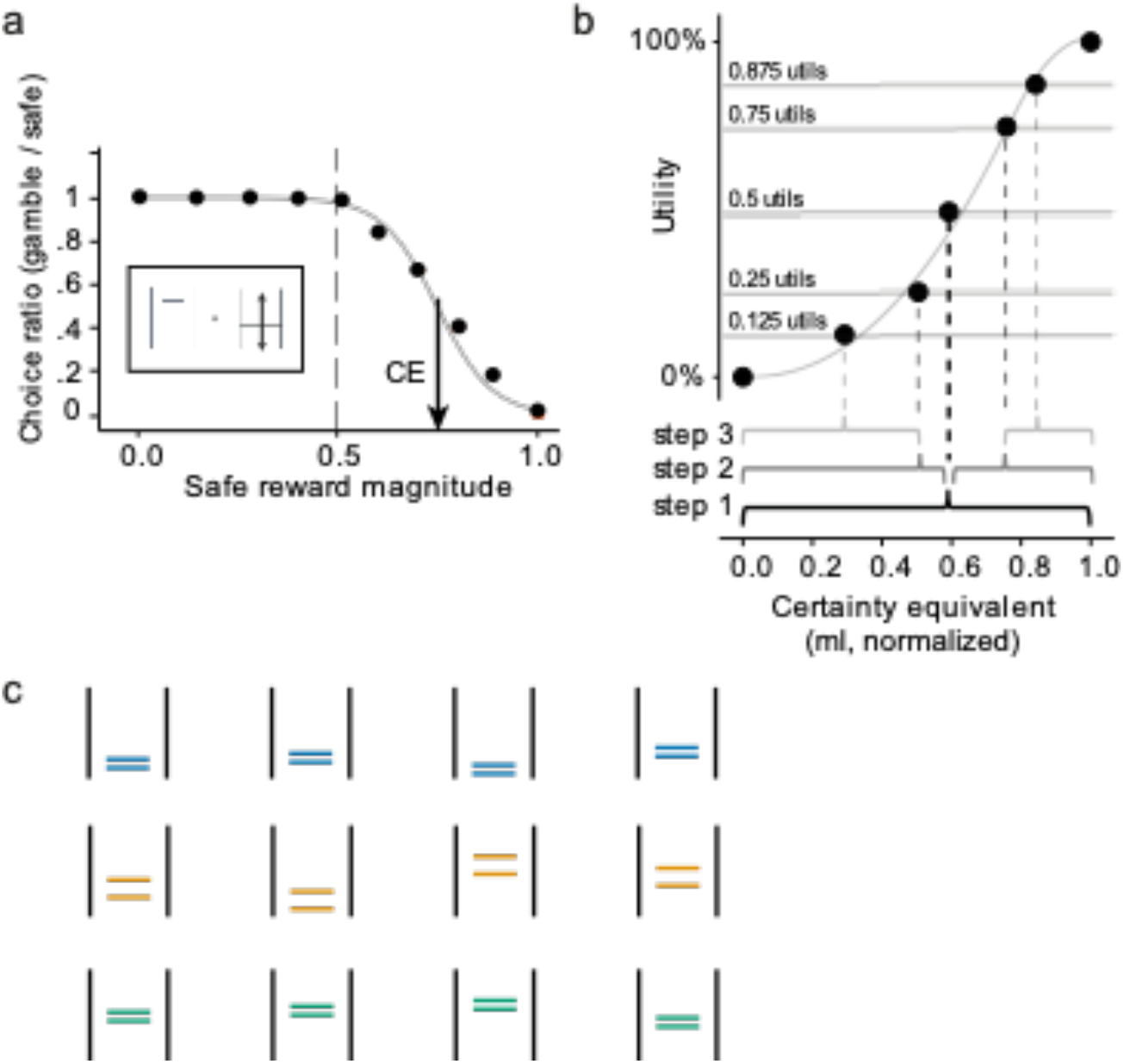
Estimating certainty equivalents and utility functions. a) Basic choice behaviour and estimation of certainty equivalents. Animals chose between a safe reward and a gamble on each trial. The safe rewards alternated pseudorandom on every trial – never going above or below the highest and lowest magnitudes tested in the daily reward distribution. Each point is a measure of choice ratio: the animal’s probability of choosing the gamble option over various safe rewards. We fit psychometric softmax functions (Eq. 1) to these choice ratios, separately for each day, and recorded the certainty equivalent (CE) of individual gambles as the safe magnitude for which the probability of either choice would be 0.5 (black arrow). The dashed vertical line indicates the expected value (EV) of the gamble represented in the box. b) Estimation of utility using the stepwise, fractile method. In step 1, the animals were presented with an equivariant gamble comprised of the maximum and minimum magnitudes in the tested reward distribution. the CE of the gamble was estimated and assigned a utility of 50%. In step 2, two new equivariant gambles were defined from the CE elicited in step 1. The CEs of these gambles were elicited and assigned a utility of 25% and 75%. Two more gambles are defined in step 3, from the CEs elicited in step 2. Their CEs were then assigned a utility of 12.5% and 87.5%. Parametric utility functions, anchored at 0 and 1, were fitted on these utility estimates (see *methods*). c) Equivariant, equiprobable gambles presented in out-of-sample validation sequences. Sets of four gambles, unique to each reward distribution, were used to validate the risk attitudes predicted by the fractile-derived utilities. The CEs of these gambles were measured (see panel a) and the difference between CEs and the specific gambles’ EVs signalled the animals’ risk attitudes: if the difference was positive, the animals were risk seeking, if the difference was negative, the animals were risk averse.

The probability of the animal choosing a safe reward over the 0.5 utility gamble (P_(ChooseSafe)_) was contingent on the safe option’s magnitude (*SafeReward_ml_* and the two free parameters, xø: the x-axis position of the curve’s inflection point, and σ: the function’s temperature. Importantly, this function’s inflection point represented the exact safe magnitude for which the animal should be indifferent between the set gamble and a given safe reward. Then we assigned utility to the lowest juice reward amount (0.0 utils) and highest juice amount (1.0 util) for the currently tested distribution (Fig. 2b). Since the animals only experienced trials set between these reward magnitudes, this constrained all utility estimates between 0 utils and 1 utils. The x0-parameter could thus be used as a direct estimate of the gamble’s CE: at choice indifference, the safe reward had the same utility as the equiprobable gamble (*p* = 0.5 each outcome) formed of these two magnitudes, which amounted to 0.5 = [0.5 * 0 utils] + [0.5 * 1 utils]). In the subsequent step, a new equiprobable gamble was set between 0 ml and the first CE’s ml value and the CE elicitation procedure was repeated (logistic fitting, Fig. 2a); their CE had a utility of 0.25 utils (1/4 of maximal utility). In the next step, two new equiprobable gambles were set between the first CE’s ml value and the maximum magnitude of the currently tested reward distribution, i. e. 0.5 ml; their CE had a utility of 0.75 utils (3/4 of maximal utility). Crucially, gamble/safe pairings for both gambles were interwoven in the same sequence – to ensure a similar spread in the presented rewards. Only sequences that contained a minimum of three different choice pairs (repeated at least 4 times) were used in the elicitation of CEs, and only the fractile sequences where at least 3 utility values could reliably be estimated were used in further analyses. The CEs assigned to each utility level, in each reward distribution, were compared via two-way ANOVA.

### Parametric estimation of utilities from aggregate and single choices

Parametric utility curves were fit onto the CE-Utility data to capture and predict an animal’s choice preferences over the entire distribution of rewards. These utility curves served as a direct signal of the animals’ risk attitude over the tested reward distribution: if the fitted utilities were convex (i. e. increasingly curving upwards) the animals had demonstrated risk seeking behaviour; if the curves were instead concave (i. e. gradually flattening), the animals had demonstrated risk aversion. Several parametric utility models were compared to ensure the most reliable utility predictions; the best-fitting functions would then be used for all further analyses. In accordance with the assumptions of the fractile method, each of these functions had to be anchored at 0% to 100% on the y-axis –– and we normalized the CEs on which they were fit to be between 0 and 1. Finally, because CEs, not utilities, were the measured data (i. e. the error was relative to the x-axis), orthogonal distance regression was used to fit each and every function (Boggs & Rogers, 2012). We fit two 1-parameter functions (U_1-Power_, U_1-Tversky_),

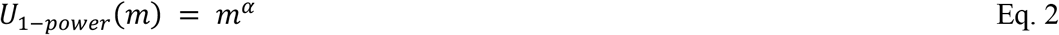

with *m* for juice magnitude (in ml) of a given reward outcome and α as power parameter of the function (if α < 1 utility function is convex, if α > 1 utility function is concave).

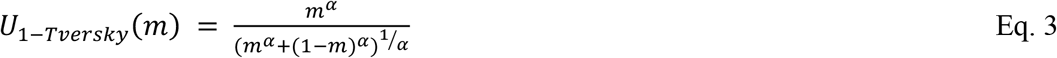

with α as temperature parameter of the function (if α > 1 utility function is S-shaped, if α < 1 utility function is inverse S-shaped).

Two 2-parameter functions U_2-Prelec_, U_2-SCDF_),

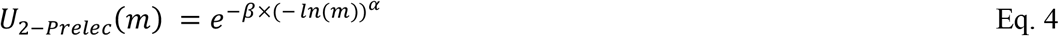

with α-parameter as temperature parameter of the function (generally, if α > 1 utility function is S-shaped, if α < 1 utility function is inverse S-shaped), and the β-parameter controls the height (or location) of the function’s inflection relative a 45° line across the x- and y-axes of the function.

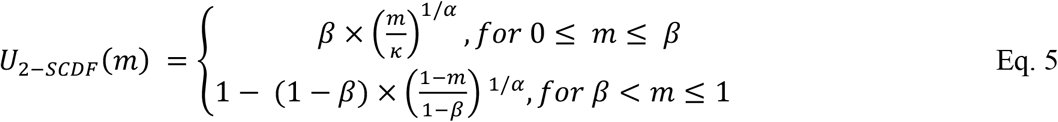

with α as the power of the function’s curvature (if α > 1 utility function is S-shaped, if α < 1 utility function is inverse S-shaped), and the β-parameter controls the x-axis position at which the function’s curvature inverts.

And one 3-parameter function (U_3-Power_)

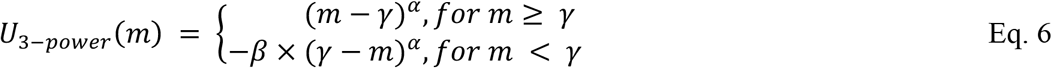

with α as the power of the function (generally, if α > 1 utility function is S-shaped, if α < 1 utility function is inverse S-shaped), the β-parameter accounts for any loss aversion.

Sets of daily Bayesian Information Criterions (BIC) were then calculated from the orthogonal residuals of each fitted model 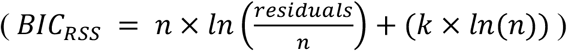. We selected the best fitting function using a one-way Friedman test followed by pairwise Wilcoxon signed-rank tests (Bonferroni-Holm corrected) and compared the estimated parameters specific to each reward distribution using a one-way MANOVA.

Since the fractile method relied on stepwise, chained measurements (where later metrics depend on earlier ones), utility functions were also estimated using a discrete choice model (DCM) fitted to single trials for comparison. By fitting a model on individual choices rather than aggregate CE sequences, we avoided the propagation of estimation errors from earlier steps onto the next and therefore reduced estimation biases for individual utility functions (Abdellaoui, 2000).

As is commonly done (McFadden, 2001; Stott, 2006), the likelihood that animals would choose the left option over the right one, given a set noise level and side bias, was modelled using a logit function:

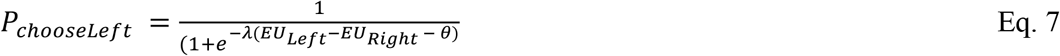

The probability of choosing the left option was, therefore, in the DCM, a function of the expected utility difference between the left and right options, the temperature (or noise) parameter, *λ*, and *θ* which captured side bias parametrically. The expected utility of each option (euleft, euright), as a function of their probability (p) and the utility function U(m), was given by the functional form:

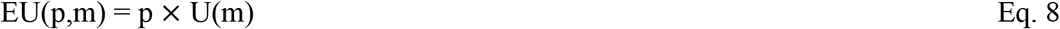

The model’s best-fitting parameters were estimated by minimizing the following cumulative loglikelihood function:

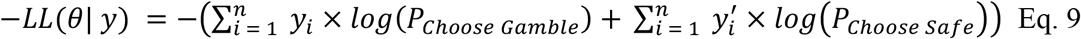

Where y and y’ indicated a left or right choice (0 or 1), respectively, for each trial i; n was the total trial number for the session.

Again, the best-fitting discrete choice model was selected via BIC comparisons, this time defined on the likelihoods (*BIC_LL_* = (*k×ln*(*n*)) − (2 × *LogLikelihood*)). The parameters estimated in each reward distribution were also using a one-way MANOVA.

### Validating utility predictions from out-of-sample certainty equivalents

To validate the predictions of the utility functions, CE measures were gathered from binary choices presented outside of the utility estimation sequences testing gambles not employed for the utility estimation. These gambles were used to corroborate the risk attitudes predicted by the fractile- or DCM-derived utilities. Two of the three animals were presented with three sets of four gambles unique to each reward distribution for which we estimated CEs. We used these 12 CEs to validate the risk-attitude predictions of the utility function estimated in each distribution. The gambles in the narrow reward distributions had a spread of 0.15 ml, while gambles in the full distribution had a spread of 0.30 ml – keeping the relative spreads equivalent across the distributions. Gamble means were also, once normalized, centred around the same relative values. In percentage points, each gamble spread over 30% of the reward distributions, and gamble was centred at a value representing 25%, 45%, 65%, or 85% of the reward distribution (Fig. 2c).

Taking the difference between the CEs of these gambles and their expected value (EV) as a proxy for risk attitude (CE – EV), the risk-attitude estimated from these CEs were compared with the predictions from the fractile-estimated and discrete-choice utility curves. If the CE – EV metric were positive, it signalled that the animals were risk seeking. If instead the measures were negative, the animals could be seen as being risk averse. Because of this, if the utility models imposed an S-shape that was unrealistic (and a consequence of the function used) the CE – EV fits would expose it right away: they would not go from risk seeking to risk averse. These measures were repositioned relative to the inflection point at which fractile- and DCM-derived utilities predicted reversal of riskattitudes (i. e. the point of risk neutrality. Linear regressions were fit to the repositioned CE – EV metrics in order to identify which of the two inflections proved most reliable in predicting out-ofsample behaviour (fractile or DCM-derived):

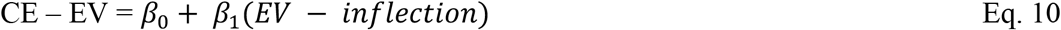

In the model, *β*_0_ Indicated the point at which CE measures became risk-neutral, and *β*_1_ Paralleled the ‘depth’ of utility’s curvature. The R^2^-value associated with both regressions was compared to see which of the two utility estimation procedures most reliably matched out-of-sample behaviour. Put simply, these regressions allowed both the validation of predicted risk-attitudes, and the selection of the better-fitting procedure.

### Defining preference adaptation metrics

Comparing the utilities estimated from choices in different reward distributionswas done in one of two ways: the first, assuming that preferences were fixed and did not adapt to the distribution of possible rewards in a task; the second, assuming that preferences fully adapted to the reward spread and magnitude of the task at hand. To test for the former, utilities estimated in narrow distributions (i. e. low- and high-distribution) were compared to the full-distribution ones. For the assumption of full adaptation, utilities were compared sequentially - looking for differences in the shape of the utilities between different distributions.

The parametric utility functions had a unique inflection point, defined as a single point where the utility function’s curvature reversed, and where the function’s first derivative was maximal. This inflection identified the precise reward magnitude for which the animals’ risk-attitude changed, and served as a good indicator for where and how the animals’ preferences would change depending on the variance and mean of the reward distribution. The inflection points elicited in different distributions were compared using a Kruskal Wallis test with Bonferroni-Holm corrected post-hoc analysis (Wilcoxon test).

Another metric, the curvature ratio (CR) was defined as the normalized area under the utility functions (the function’s area divided by the total area in each distribution). The CR provided a direct, normalized metric of the convexity/concavity interplay of daily utility estimates – reflecting overall risk attitude to a greater degree than inflection points. A linear utility function would have a CR of 0.5, as would perfectly symmetric S- or inverse S-shaped utilities. A CR above 0.5 indicated that the functions fell above the diagonal and predicted risk averse choices; conversely, a CR under 0.5 reflected more risk seeking choices. The CRs measured in the different distributions were also compared using a Kruskal Wallis test followed by pairwise Wilcoxon rank sum comparisons (Bonferroni-Holm corrected).

A final series of metrics, defined as adaptation coefficients, allowed for the quantification of relative changes in CRs. Between utilities that had been estimated in consecutive reward distributions. A sequential adaptation coefficient (SAC) was calculated as:

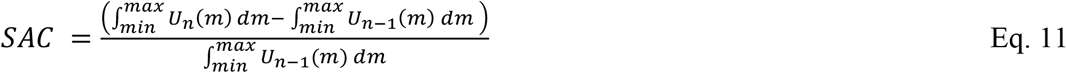

And it captured changes in the median utility of a given reward distribution n (*U_n_*(*m*)), where m represented every reward between the minimum and maximum rewards in the tested distribution, relative to the median utility function in distribution n-1 (*U*_*n*−1_)(*m*)). Since all parametric functions were defined from 0 to 1, comparing the area under each curve gave us a direct measurement of the difference between the utilities that captured preferences in consecutive reward distributions.

A second coefficient, the general adaptation coefficient (GAC), compared the utility of low- and high-reward distributions to the utility estimated from a animal’s full reward distribution. The GAC placed the narrow-distribution utilities (i. e. the low and high distribution ones) relative to the shape of the full-distribution’s utility function. That is, a GAC of 0 would indicate that the narrow-distribution utilities are but segments of a fixed full-distribution one, whereas a GAC of 1 suggested that utilities kept a similar form but fully shifted to represent preferences in the new distribution. For any GAC where 0 < GAC < 1, utilities had partially adapted. To calculate this, narrow distribution utilities were rescaled to map onto the full distribution ones: the maximum value of the low-distribution became the utility value of the full-distribution utility at 0.5 ml, and the utility value of the full-distribution utility at 0.5 ml became the minimum value of the high-distribution. Then, the median utility of the full distribution (*U*_Full_) was rescaled (into *U_adapt_*) to match the domain and image of narrow distribution utilities (U_Low-distribution_ and U_Hugh-distribution_). The GAC was given by

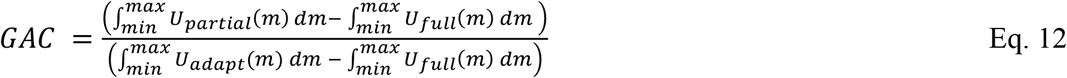

Where *min* and *max* are the minimum and maximum reward magnitudes in a narrow distribution condition. A GAC of 1 signalled full adaptation while a GAC of 0 indicated that no adaptation had taken place. Crucially, the GAC metric took no account of the order in which reward distributions were tested; it relied instead on full-distribution utility function as a comparison template.

## Results

### Experimental design

In order to investigate the adaptation of utility functions to different reward distributions, macaque monkeys were presented with sequences of binary choices while reward distributions were kept constant over consecutive days and then suddenly changed. Thus, without other task changes, the animals experienced periods of relatively low reward reward magnitudes, periods of relatively high magnitudes, and periods with a mix of both (Figs. 1c; 3). On each day the animals were presented with either a utility estimation sequence, an equivariant gamble sequence (out-of-sample validation), or both.

In utility estimation sequences, utility measurements were derived from the choices that animals made between sets of gambles and safe rewards. Using the fractile method (see *Methods*), utilities were derived from the certainty equivalents (CEs) of specific sets of binary, equiprobable gambles (*p* = 0.5 each outcome; the magnitude of safe reward that was subjectively equivalent to the gamble). In validation sequences, the animals’ risk preferences were measured directly using the CEs of out-of-sample binary, equiprobable gambles. These measurements were then used to confirm the utilities estimated in elicitation sequences.

For each reward distribution, sets of daily utilities were estimated using the fractile method. The way reward magnitudes (CEs) mapped onto these utilities (once normalized to the minimum and maximum rewards in a distribution) could then be compared within and between the different rewards distributions. To do so, and because utilities were defined from 0% to 100% regardless of their distribution, the CEs were normalized relative the maximum and minimum magnitudes in the appropriate reward distribution (Fig. 3). As expected, higher utility values mapped onto higher reward magnitudes (higher CEs), but the way in which they did so differed markedly depending on the current distribution. The same utility levels (12.5%, 25%, 50%, 75% and 87.5%) in different reward distributions did not map onto the same relative magnitudes (i. e. normalized CEs). We confirmed this statistically using a two-way ANOVA with the main factors being the utility level tied to individuals CEs and the reward distribution from which they had originated. The ANOVA confirmed that there was a significant main effect of utility level on the value of the estimated CEs (Monkey A: F(4,295) = 64.301, p = 4.812× 10^−39^; Monkey B, F(4,192) = 50.51, p = 4.107× 10^−39^; Monkey C: F(4, 295) = 609.547, p = 3.254× 10^−141^; The distribution in which utility-specific CEs had been estimated also had a significant main effect on the value of the estimated CEs (Monkey A: F(2,295) = 356.415, p = 1.991× 10^−79^; Monkey B, F(2,192) = 8.994, p = 0.003× 10^−3^; Monkey C: F(1, 295) = 16.204, p = 7.235× 10^−5^). Together, these corroborated what we could see graphically (Fig. 3): higher CEs correlated with higher utilities in all distributions, but these CEs were all relatively lower once a shift from low- to full- or high-distribution had occurred. Supporting the two other main effects, there was a significant interaction effect of utility level and distribution on the estimated CEs, in two of the three animals (Monkey A: F(8,295) = 1.156, p = 0.326; Monkey B, F(8,192) = 5.217, p = 1.829× 10^−6^; Monkey C: F(4, 295) = 8.488, p = 1.707× 10^6^). That is, the steepness of the utility-CE pairings changed between the different reward distributions – rather than simply shifting and recalling, utilities in different distributions seemed to follow different patterns.

**Figure 3.**
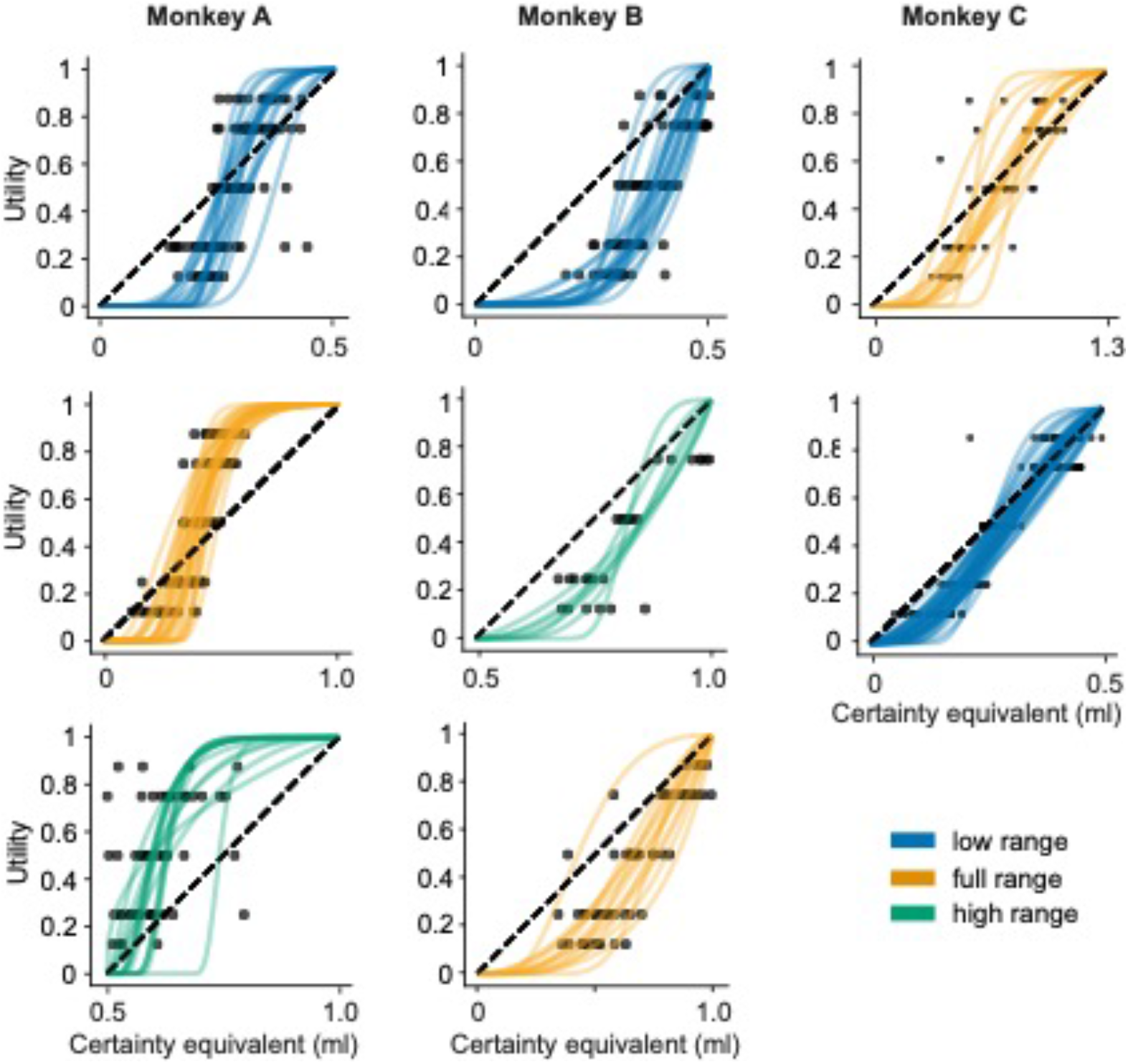
Utility functions elicited from daily fractile procedures. Order of distributions tested is captured vertically. Black dots represent CE-utility pairings elicited in individual experimental sessions using the fractile method; coloured lines are parametric fits (*U*_2−*prelec*_) to daily CE estimates (blue, low narrow distribution; yellow, full distribution; green, high narrow-distribution. Utility fits for Monkey A, from top to bottom, represent 20 days, 26 days, and 15 days. For Monkey B, we have 23 days, 7 days, and 13 days. Finally, Monkey C has a total of 13 days for the top panel, and 43 days for the lower one. In all cases, convexity of the functional fit signals risk seeking behaviour, concavity signals risk aversion.

### S-shaped utilities best fit choices

Parametric utility functions were fitted to the daily utility measurements to better compare and understand the relationship between the utilities estimated in each distribution. To do so, several different functional forms of utility were first compared; the most reliable function was then used for all further analyses. Power functions are commonly used to model utility functions. We therefore fit a 1-parameter power (U_1-Power_), 2-parameter CDF of a two-sided power (U_2-SCDF_), and a 3-parameter anchored power functions (U_3-Power_) to the animal’s CE-utility pairings. In addition to power-type functions, we looked at functions typically reserved for probability distortion modelling (Ferrari-Toniolo et al., 2019; Stott, 2006): the 1-parameter Tversky function (U_1-Tversky_), and the 2-parameter Prelec (U_2-Prelec_) – two functions that could readily take on the s-shape prescribed by PT. All functions mapped reward magnitudes onto utility values from 0 to 1 (i. e. 0% to 100% of normalized utilities), and all but the 1-parameter power function could capture risk seeking and risk averse behaviour, as well as any inversion in the animals’ risk attitudes within a reward distribution.

Because of the fractile method’s reliance on aggregate, chained datapoints (Farquhar, 1984; Machina, 1987), utility functions were also fit using a discrete choice model (DCM) applied to individual, rather than aggregate, choices (Eq. 7). In line with the fractile-derived utilities, and because previous experiments with the same animals had identified negligible probability distortions for p = 0.5 (Stauffer et al., 2015), choices in the model were then predicted based on the choices’ expected utilities (probabilities were treated as objective). The parameters that best described individual choices in each model were estimated through maximizing the cumulative log likelihoods of the DCMs defined on individual experimental sessions (Eq. 9; see methods).

To select the utility function that best described both the CEs and individual choices, we used the Bayesian information criteria (BIC) from all fitted models; the model with the lowest median BIC would thus represent the best fitting model. Of the five tested utility functions, the 2-parameter Prelec proved most reliable in fitting both forms of data (Fig. 4a,b). Though the model is normally reserved for probability distortion models, it presented the lowest BICRSS score as derived from the residuals of fractile-derived utilities (significantly so, Friedman test; Monkey A: F_r_(4,240) = 177.154, p = 3.046× 10^−37^; Monkey B: F_r_(4,168) = 140.780, p = 1.903× 10^−29^; Monkey C: F_r_(4,220) = 120.800, p = 3.604× 10^−25^), and the lowest BIC_LL_ score as derived from the log likelihoods of the discrete choice fits in 2 of 3 monkeys (Friedman test; Monkey A: F_r_(4,240) = 219.091, p = 2.327× 10^−45^; Monkey B: F_r_(4,168) = 186.469, p = 2.221× 10^−38^; Monkey C: F_r_(4,220) = 180.020, p = 5.298× 10^−37^). In Monkey A, the BIC_LL_ of the 2-parameter CDF of the two-sided power distribution and the 2-parameter Prelec proved statistically indistinguishable. From these BICRSS and BIC_LL_ measures, and because the behavioural predictions from each fitting method generally agreed (Fig. 4c), we selected the 2-parameter Prelec function for all further analyses.

**Figure 4.**
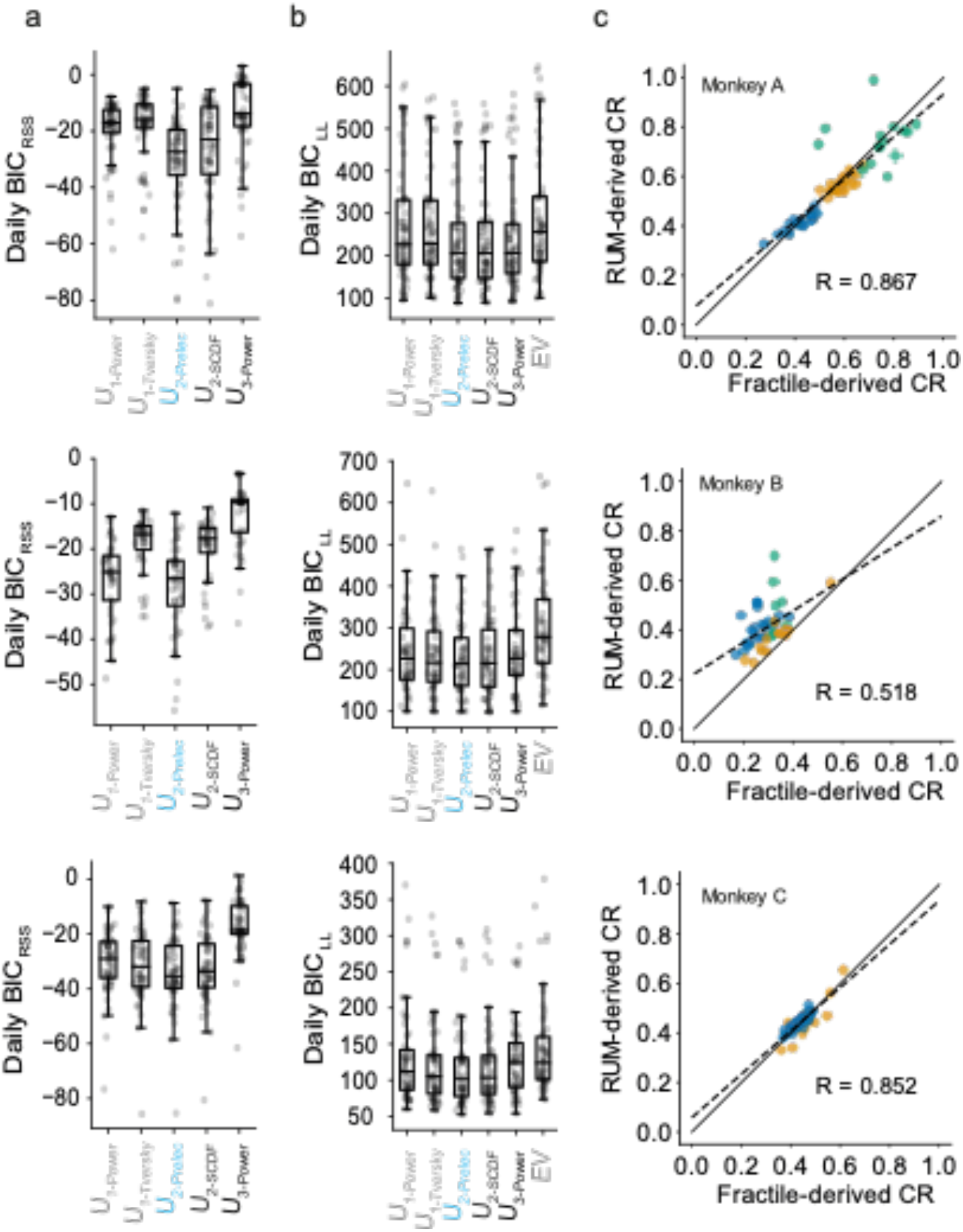
Model comparisons within and across fitting procedures. a) Model selection for fractile-derived utilities. We calculated daily Bayesian information Criterions for each utility function using the orthogonal residuals on each fit (BIC_RSS_). Lower BIC_RSS_ scores indicated a better fit to the CE-utility pairings, and the 2-parameter Prelec model that was used throughout this study appears in blue (U_2-Prelec_). b) Model selection for discrete choice utilities. We again calculated daily BIC scores for each utility function, this time using the log-likelihoods estimated to fit each discrete choice models (BIC_LL_). Lower BIC_LL_ scores indicated better fits between the discrete choice model (DCM) predictions and individual measured choices pairings. Again, the 2-parameter Prelec model that was used throughout this study appears in blue (U_2-Prelec_), and, in contrast to the fractile-fits, we also compared the various DCMs to predictions based on expected value (seeing if noise alone could explain choices). c) Curvature ratios (CRs) from each fitting procedure correlate. We calculated CRs as the area under the curve of each utility function. Each point represents the CRs from fractile-derived utilities (x-axis) and DCM-derived utilities (y-axis); their colour captures the reward distribution from which they estimated (blue: low-distribution, green: high-distribution; yellow: full-distribution). Significant positive correlations between the fractile-derived CRs and DCM-derived CRs were found in each of the three animals, and we only observed clear differences between the two procedures in Monkey B.

### Risk preferences adapt to novel reward distributions

Each fitted utility function provided a pair of parameters that could be compared to those elicited in the same or different reward distributions. The curvature of these utility functions served as a direct indicator of the animal’s risk attitude for any given magnitude. Convexity reflected risk seeking behaviour; concavity signalled risk aversion. From these parametric functions, three predictions could be made: utilities would either i) fully adapt to the novel reward distributions, ii) not adapt and remain constant (i. e. different parts of the same curve), or iii) utilities would partially adapt in a way that did not solely rely on the current reward distribution. To test for these predictions, further analyses were split into two sets of hypotheses. One set looked at utilities under the assumption that no adaptation had occurred, the other assumed full utility adaptation between each of the reward distributions. In the case of the no-adaptation assumption, the predictions from utilities on identical reward magnitudes in the narrow distribution and full distribution were compared (Fig. 5a). For the full adaptation assumption, the utilities from sequential reward distributions were normalized and compared, looking at any differences with the previous distribution’s pattern of risk attitude (Fig. 5b, c). If neither assumption proved accurate, then the assumption would be that neither full nor no adaptation had taken place – that is, preferences would have partially adapted.

**Figure 5.**
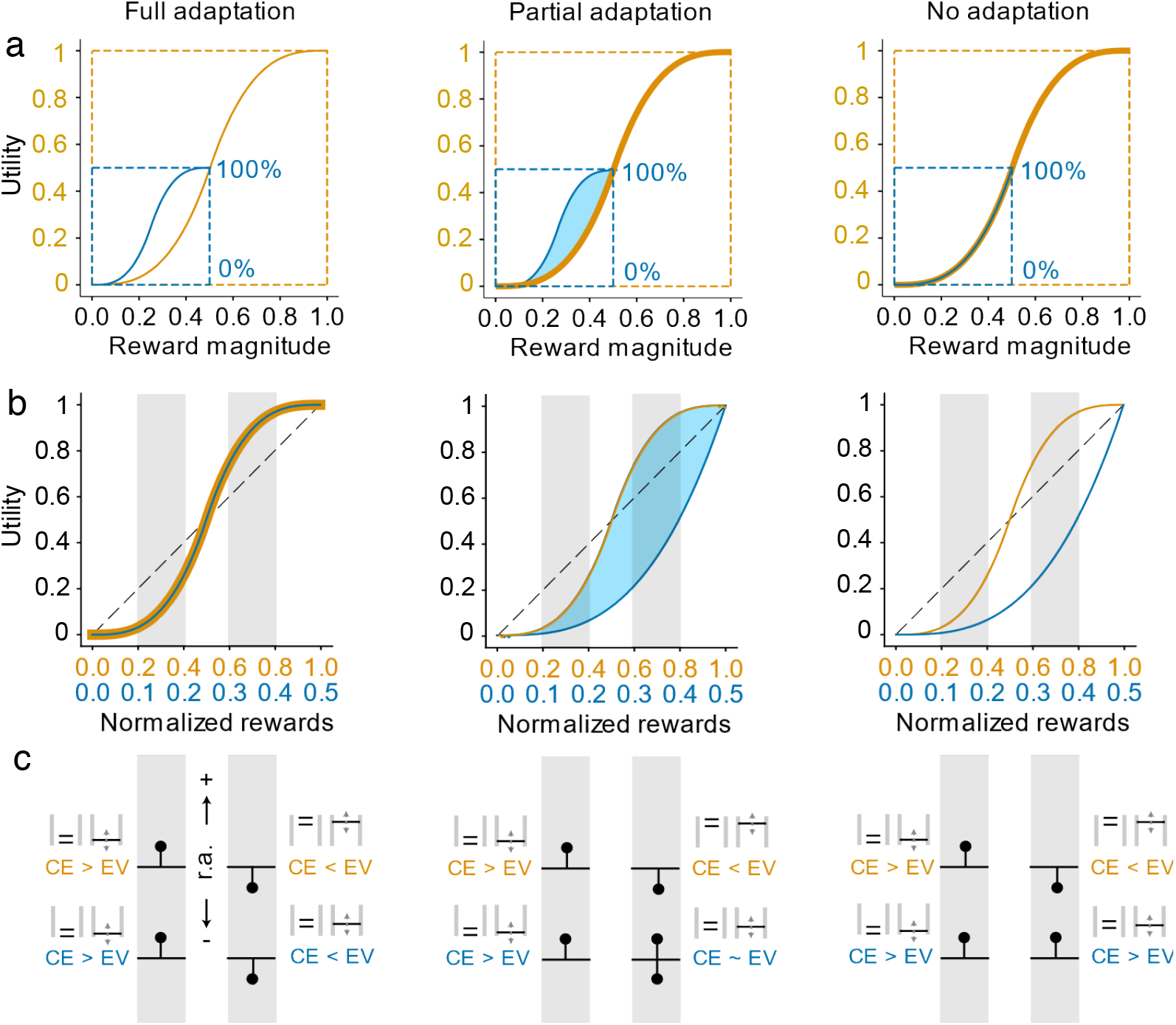
Schematic representation of full-, partial-, and non-adapting utilities estimated in low- and full-distributions of rewards. a) Scaled, identical utility functions in different reward distributions: the utility value of a 0.5 ml reward in the small distribution (blue curve, 100% utility) is scaled to the utility value of 0.5 ml reward in the large distribution (yellow curve). From left to right, utilities reshape assuming full-, partial-, and no adaptation. The three possibilities differ mostly in terms of the risk-attitudes exhibited for rewards between 0 ml and 0.5 ml – under full adaptation they should differ, under no adaptation they should not. b) Utilities normalised according to the reward distribution from which they were estimated. Utilities are set on the same scale by normalizing across the domains of each function. Curves should overlap if utilities adapt fully (left) and fail to do so if there is no adaptation (right). If functions fail to adapt the low distribution utility is predicted to be identical to the first half of the full distribution utility curve. c) Predicting the direction of risk attitudes (r.a.) from utilities. For an equiprobable gamble made up of the two outcomes that fall at the edges of each grey shaded area, the horizontal black line depicts the expected value (EV) and the black dot above or below signals the direction in which we expect the certainty equivalent (CE). A black dot above the horizontal line signals risk seeking behaviour (or positive r.a.) and a CE of higher value than the EV, and a dot below the line signals risk averse behaviour (negative r.a.). From left to right we again have predictions of r.a. given full-, partial-, or non-adaptive preferences.

Starting with fractile-derived utilities, comparing the functional parameters elicited in the different reward distributions provided us with a stringent test regarding the full adaptation assumption. In the 2-parameter Prelec function, the α-parameter represented the temperature of the function, while the β-parameter captured the relative height of the curve. If these were identical across conditions, similar patterns of utility reflected preferences regardless of unique reward magnitudes in the different reward distributions. One-way MANOVA analysis on the log-transformed parameters confirmed that this was not the case: there was a significant effect of reward distribution on the parameters elicited in each condition, for all animals (Monkey A: F(2,59) = 34.913, Wilks’s λ = 0.454, p = 1.116× 10^−10^; Monkey B, F(2,41) = 13.695, Wilks’s λ = 0.594, p = 2.946× 10^−5^; Monkey C: F(1, 54) = 9.381, Wilks’s λ = 0.739, p = 3.252× 10^−4^). Specifically, there was a significant difference between Monkey A and B’s β-, or height-, parameters (Monkey A: F(2,59) = 67.301, p = 2.447× 10^−11^; Monkey B, F(2,41) = 13.695, p = 2.946× 10^−05^; Monkey C: F(2,54) = 1.120, p = 0.290), as well as a significant difference in Monkey C’s α-, or temperature-, parameters (Monkey A: F(2,59) = 0.434, p = 0.513; Monkey B, F(2,41) = 2.583, p = 0.116; Monkey C: F(2,54) = 18.858, p = 6.236× 10^−5^). The utilities, in terms of parameters, differed depending on the distribution from which they were elicited (Fig. 6).

**Figure 6.**
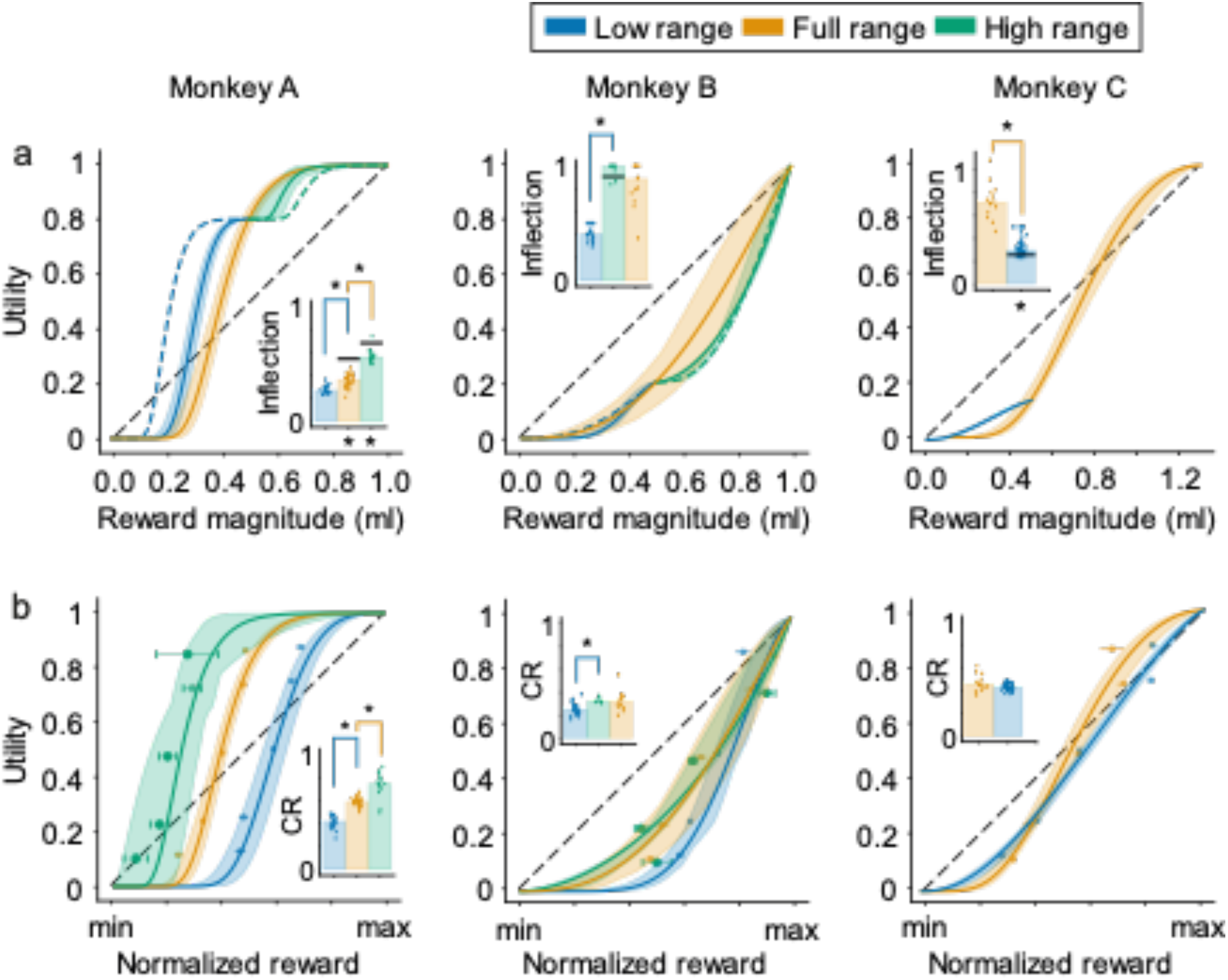
Fractile-derived utilities reflect adaption to different reward distributions. a) Scaled utilities estimated from fractile-derived CE-utility pairings. Each curve represents the median of daily, distribution-specific parameter estimates; 95% Confidence intervals were estimated via boostrapping said parameters (random sampling with replacement, n=10000). Dotted blue lines represent predictions full-distribution utilities predicted to fully-adapt to low-distributions. The dotted green lines represent similar full-adaptation predictions in the high distribution. Bar graphs represent the median inflection point, i.e., the reward magntiude at which the curve goes from convex to concave – points are daily inflection points. Upper asterisks (*) indicate differences between daily inflection estimates in two sequential distributions (Wilcoxon rank sum test, p < 0.05); Lower asterisks (*) indicate significant difference between the median predicted inflection from the previous tested distribution and the true inflection estimates of the next distribution (Wilcoxon rank sum, p < 0.05). b) Normalized utilities estimated from fractile-derived CE-utility pairings. Each curve is the median of daily, distribution-specific parameter estimates normalized according to the minimum and maximum rewards in the tested distribution. Again, 95% confidence intervals were estimated via boostrapping. Points represent mean normalized certainty equivalents ± SEMs for each of the tested distribution. Bar graphs representmedian curvature ratios (CRs) for each distribution; the relative concavity of each utility (concave > 0.5; convex < 0.5) – individual points are daily CRs. Upper asterisks (*) indicate significant differences between CRs estimated in sequential distributions (Wilcoxon rank sum, p < 0.05). For each panel, blue comes from low-distribution utilities, yellow from full-distribution, and green from high-distribution.

To explore how these parametric differences influenced utility patterns in a way that was directly comparable between conditions, we compared the position of each utility function’s inflection points – the reward magnitude at which the behaviour predicted by the utility function flipped from risk seeking to risk averse (or risk averse to risk -avskseeking depending on the temperature of the utility function). The inflection crudely summarized choice predictions with a single metric – one that had been previously used to signal animals’ ‘reference-points’ (Chen et al., 2006; Lakshminarayanan et al., 2011). Importantly, since this metric was tied to CE values; one could easily observe if inflection points fell on similar magnitudes depending on the distribution in which it had been measured (Fig. 5a).

From these inflection points, the assumption of no adaptation was tested by comparing both within and across-distribution inflections. If no adaptation had occurred, the inflections would be the same within and across the different reward distributions. Testing for the former, i.e. Within distribution differences in inflection points, no significant pattern of change could be identified – at least for Monkeys A and B (linear regression analysis, Monkey A: p_full-distribution_ = 0.160, p_high-distribution_ = 0.472; Monkey B: p_full-distribution_ = 0.270, p_high-distribution_ = 0.714; Monkey C: p_low-distribution_ = 0.009). And since Monkey C’s low distribution had been tested over a year after changing distributions – the fact that a significant positive slope was identified (the inflection slowly went up in value over the days of testing) did little to indicate distribution-swap adaptation. Moving from within distribution to between distribution analyses, there were significant differences between the distribution-specific inflections for all monkeys (Kruskal Wallis test; Monkey A: H(2,58) = 44.281, p = 2.424× 10^−10^; Monkey B: H(2,40) = 27.973, p = 8.429× 10^−7^; Monkey C: H(1,54) = 28.397, p = 9.881× 10^−8^), which translated into significant pairwise differences (Wilcoxon rank sum) for all but Monkey B’s high and full distribution inflection points (Fig. 6a). Simply put, the inflection points fell on different reward magnitudes for each of the reward distributions. If preferences had truly been non-adaptive, no significant difference across any of the conditions would have been observed.

Since none of the results corroborated the no-adaptation hypothesis, the next step was to test for full adaptation. Rather than comparing the absolute position of the utilities’ inflection points, testing for full adaptation required predicting where inflection points from a past distribution would map onto the next distribution: the assumption being that if the same utility function simply shifted to a new distribution (i. e. fully adapted), the relative position of the inflection should be the same. An inflection at 0.3 ml in the low distribution, for example, would be placed at 0.15 ml in the full distribution, and vice versa. However, since an inflection of 0.3 in the low distribution would result in a negative magnitude when compared with the high distribution, inflections < minimum reward were set at the minimum, and inflections > maximum reward were set to the maximum. There were significant differences between all consecutive comparisons in Monkeys A and C, and none for Monkey B (Fig. 6a; Wilcoxon rank sum test). From a full adaptation perspective, this suggested that, while Monkeys A and C had not fully shifted their reference to accommodate the new distributions, Monkey B’s preferences seemed to follow the same relative pattern across all rewards distributions.

From the inflection points, the picture that emerged was one of (at least) partial adaptation. That is, the significant differences between the inflection points corroborated neither the idea of fully- or non-adaptative preferences. Nevertheless, because inflection points carried no information about the risk attitude that followed or preceded them, the inflection points could be similar even if the animals’ choices were not. To counter this, the previous comparisons were repeated using the area under each utility curve – a direct indicator of the convexity/concavity patterns within single utilities. Rather than representing a single point, the area under each curve reflected the order and intensity of risk seeking or risk averse behaviour throughout the reward distribution. Hereafter defined as curvature ratios (CRs, see methods), the areas calculated in each distribution were compared through Kruskal Wallis test (followed by pairwise Wilcoxon rank sum post-hoc tests). The results validated the earlier findings from the inflection comparisons: sequentially, there were significant differences across distributions for Monkey A and B (Monkey A: H(2,58) = 27.973, p = 8.428× 10^−7^; Monkey B: H(2,40) = 12.124, p = 0.002), but there were no statistical differences between monkey C’s CRs across conditions (Fig. 6b; H(1,54) = 1.872, p = 0.171). In essence, while the risk attitudes that Monkeys A and B exhibited differed between reward distributions, Monkey C seemed to exhibit relatively similar behaviour in the two distributions (albeit with a slightly different inflection).

To validate these fractile-based comparisons, we repeated the full/no-adaptation analyses using the DCM-derived utilities. Both the inflection points and the CRs of Monkey A reliably mimicked earlier findings: significant differences between the distributions meant inflection points were somewhat adaptive (Fig. 7a; Kruskal Wallis, H(2,58)= 44.504, p = 2.167 x 10^−10^), but differences in sequential predictions also meant that inflections were not fully-adaptive (Fig. 6a; Wilcoxon rank sum, Z(45)_full-distribution_: -5.761, p = 8.351 x 10^−9^; Z(40)_high-distribution_: -4.790, p = 1.661 x 10^−6^). Corroborating the latter, CRs were again found to be significantly different across all distribution conditions (Fig. 7b; H(2,58) = -51.342, p = 7.100x10^−12^). For Monkey B, the DCM-derived inflection points also behaved like those estimated from fractile utilities: there were significant differences between all but the high and full-distributions (H(2,40) = 31.103, p = 1.762 x 10^−7^), suggesting that inflections were not fixed, which was validated by the finding that there were no significant differences between all consecutive predictions (Z(29)_high-distribution_ = 1.103, p =0.270; Z(20)_full-distribution_ = 1.941, p = 0.052). In terms of curvature ratios, i.e. Test of no adaptation, there again was a difference between the CRs gathered in different reward distributions (H(2,40) = 7.470, p = 0.024), but this time none of the post-hoc pairwise comparisons reached significance once corrected for multiple comparisons (Wilcoxon rank sum; Fig. 6b). This meant that Monkey B’s preferences were much closer to being fully adaptive than not. Finally, Monkey C’s results, like Monkey A, were consistent across elicitation methods. Inflection points were significantly different between the two distributions tested (H(1,54) = 30.524, p = 3.297 x 10^−8^), consecutive inflection predictions were also significantly different (Z(55)_low-distribution_ = 2.076, p = 0.03), and CRs were not (Z(55)_low-distribution_ = 0.0178, p = 0.897). Inflections differed, but risk attitudes did not.

**Figure 7.**
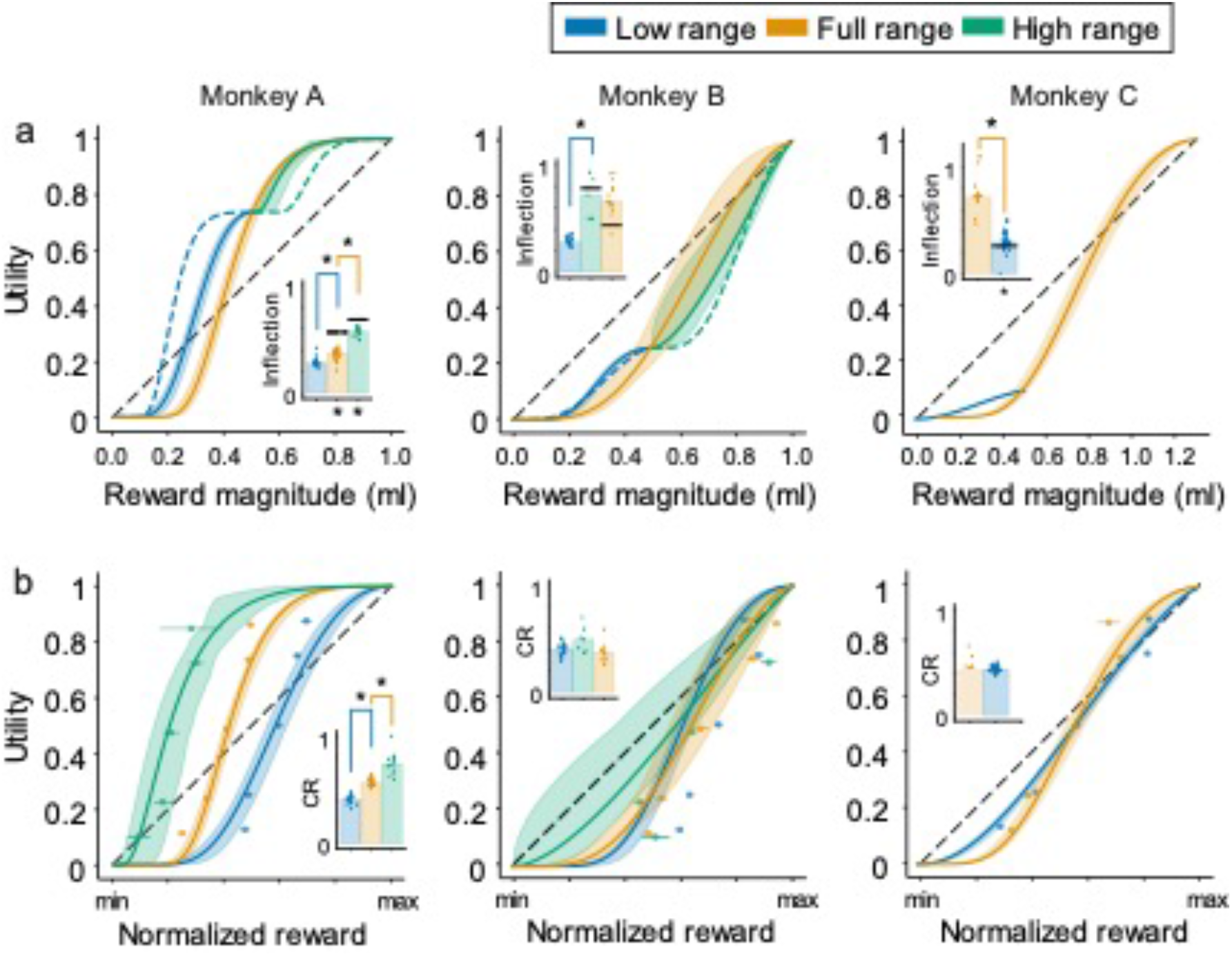
Discrete choice utilities reflect partial adaption to reward distributions. a) Scaled utilities estimated from discrete choice models (DCM). Each curve represents the median of daily, distribution-specific parameter estimates; 95% Confidence intervals were estimated via boostrapping said parameters (random sampling with replacement, n=10000). Dotted blue lines represent predictions full-distribution utilities predicted to fully-adapt to low-distributions. The dotted green lines represent similar full-adaptation predictions in the high distribution. Bar graphs represent the median inflection point, i.e., the reward magntiude at which the curve goes from convex to concave – points are daily inflection points. Upper asterisks (*) indicate differences between daily inflection estimates in two sequential distributions (Wilcoxon rank sum test); Lower asterisks (*) indicate significant difference between the median predicted inflection from the previous tested distribution and the true inflection estimates of the next distribution (Wilcoxon rank sum). b) Normalized utilities estimated from DCMs. Each curve is the median of daily, distribution-specific parameter estimates normalized according to the minimum and maximum rewards in the tested distribution. Again, 95% confidence intervals were estimated via boostrapping (random sampling with replacement, n=10000). Points represent mean normalized certainty equivalents ± SEMs for each of the tested distribution. Bar graphs represent median curvature ratios (CRs) for each distribution; the relative concavity of each utility (concave > 0.5; convex < 0.5) – individual points are daily CRs. Upper asterisks (*) indicate significant differences between CRs estimated in sequential distributions (Wilcoxon rank sum). For each panel, blue comes from low-distribution utilities, yellow from full-distribution, and green from high-distribution.

Taken together, these results suggest that while no animal (except perhaps Monkey B) demonstrated full adaptation, some form of partial adaptation had occurred across every distribution in every animal. More specifically, while not fully adapted, Monkey A and C’s utilities did shift following changes in the task’s reward statistics. Their inflection points moved, but not to the degree predicted by a full shift of the previous distribution’s inflections. Where the two animals differed, however, was in the fact that Monkey C had maintained a very similar CR across conditions – likely due to the time elapsed between the different tests. Monkey B, on the other hand, maintained the relative inflection predicted across conditions and a similar (though different in fractile-estimates) utility shape.

### Predicting distribution-specific preferences from adapting utilities

While the fractile- and DCM-fits generally agreed on the inflection of utility functions (Figs. 6a; 7a), variations in parameter estimates and concavity/convexity patterns (particularly in Monkey B; see Table. 1) highlighted the need to select the most reliable fitting procedure if quantification of adaptation was the goal.

**Table 1.**
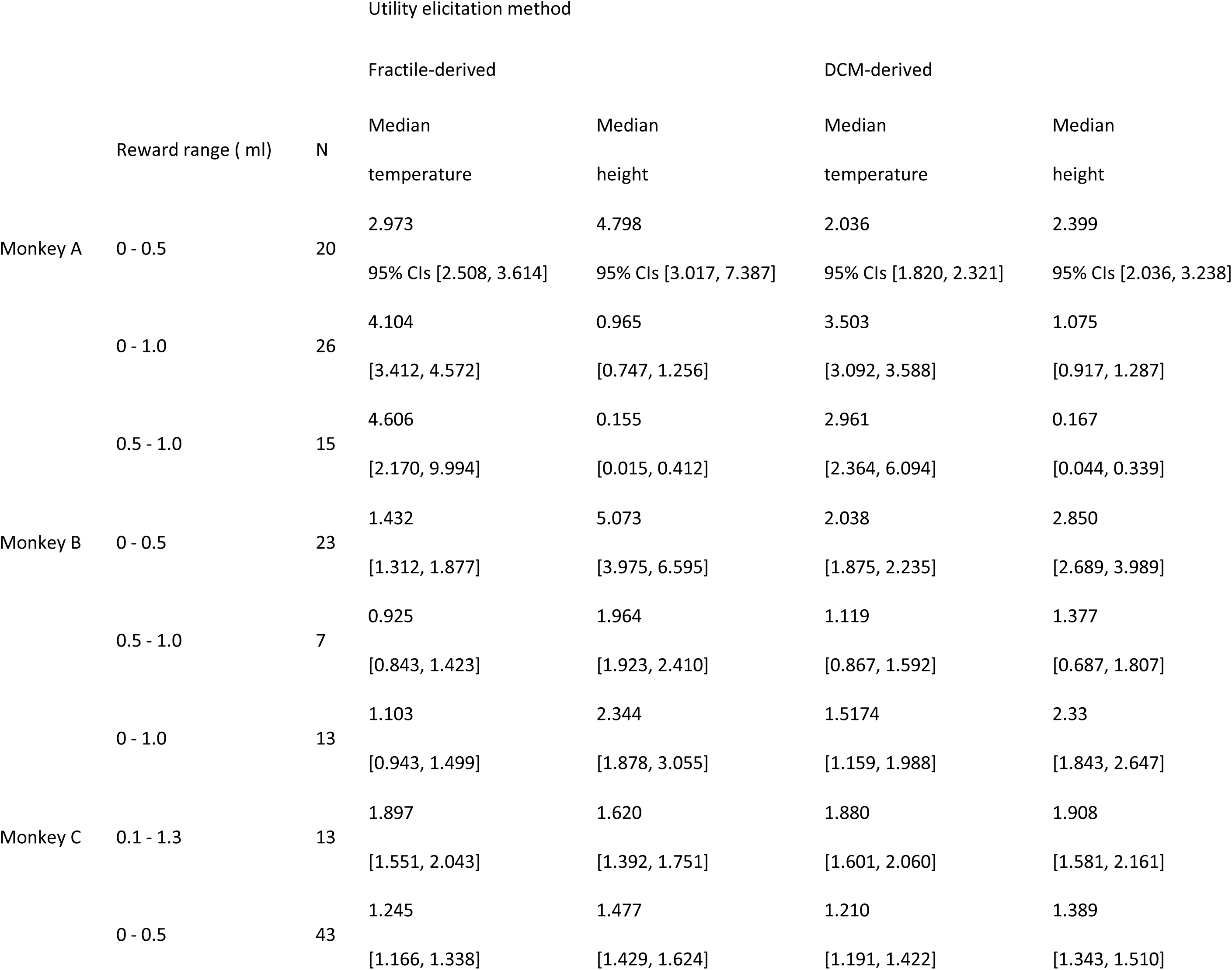
Height and temperature parameters from fractile-derived and DCM-derived utilities

To address this concern, we compared the risk attitudes predicted by the utilities of each method to real risk attitudes measured in different, out-of-sample choices (i.e. validation sequences). The CEs of equiprobable and equivariant gambles were recorded in each of the reward distributions, and the differences between these CEs and the gambles’ EVs (CE – EV) were used to indicate the animals’ risk attitudes. Every gamble had a magnitude spread equivalent to 30% of the respective reward distribution, and their EV were anchored at 25%, 45%, 65%, and 85% of the testing distribution’s magnitudes (Fig. 2c). If the difference between a gamble’s CE and its EV (CE - EV) was positive, it reflected a risk seeking attitude towards the gamble; if, on the other hand, this value was negative, the animal was said to be risk averse. These ‘validation’ measurements were gathered in two of our three animals (namely Monkeys A and B).

The CE - EV attitude predictions were compared to the risk attitude predictions from the fractile and DCM utility estimates. If the S-shaped pattern of utilities elicited for each animal were accurate, choices involving magnitudes that fell below the utility’s inflection point should have been risk-prone, while choices above it should have been risk averse (also validating the S-shape utilities as more than just an effect of the Prelec functional form). We found that this was indeed the case and that CEs in all distributions reflected both risk seeking and risk averse behaviour depending on the relative magnitudes involved (Fig. 8a). Then, to identify the best-fitting utility estimation procedure, the CE – EV values were regressed onto the gamble’s relative distance from the median inflections in each distribution (the distance in EV terms; see Eq. 10). In both animals, positioning CE – EV values relative the DCM-derived inflection resulted in a better regression fit than using the fractile-derived inflections (Fig. 8b, c) – the DCM-derived utilities were therefore chosen for further quantification as they represented a more accurate depiction of the animals’ behaviour.

**Figure 8.**
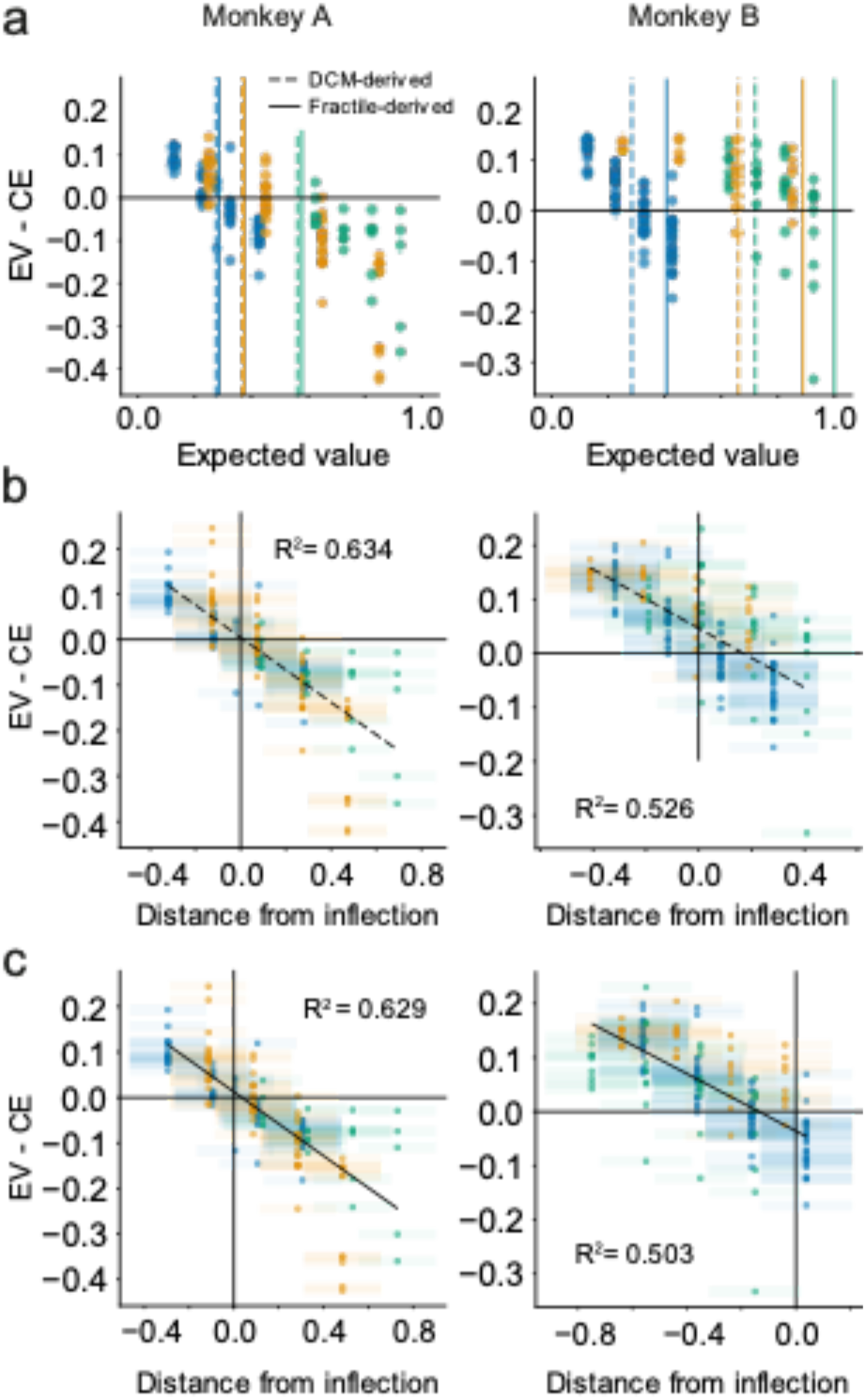
Discrete choice utilities better predict out-of-sample risk attitudes. a) Differences between the certainty equivalent (CEs) and expected value (EV) of out-of-sample, equivariant gambles reflects the risk attitudes predicted by utilities. Each point represents a CE – EV measure from individual CE estimates. For CE-EV measures above 0 reflect risk seeking behaviour, points below 0 reflect risk averse behaviour. The transition from risk seeking to risk averse behaviour should correlate with the inflection points predicted from utility functions: full lines represent the median inflection as predicted from daily fractile-derived utilities; dotted lines represent the median inflection from DCM-derived utilities. b) Discrete choice (DCM) derived inflections (better) predict risk attitudes as measured in out-of-sample gambles. CE – EV metrics positioned as a function of a gamble’s EV position relative the median fractile-derived inflection for each distribution. The x-axis captures the relative difference between the distribution’s inflection point (in ml) and a gamble’s EV (in ml). Dotted lines represent linear regression lines across all CE – EV measurements (Monkey A: p=1.77 x 10-35; Monkey B: p=1.90 x 10-31). c) Fractile-derived inflections predict risk attitudes as measured in out-of-sample gambles. CE – EV metrics positioned as a function of a gamble’s EV position relative the median fractile-derived inflection of each distribution. The x-axis captures the relative difference between the distribution’s inflection point (in ml) and a gamble’s EV (in ml). Dotted lines represent linear regression lines across all CE – EV measurements (Monkey A: p=5.43 x 10-35; Monkey B: p=1.43 x 10-29).

### Partial adaptation to reward distribution shapes risk preferences

Two final metrics served to quantify the degree to which each animal’s DCM-utilities had adapted between the different reward distributions: a sequential adaptation coefficient (or SAC; Eq. 11) and a general adaptation coefficient (GAC; Eq. 12). The SAC served to quantify how the utilities adapted sequentially as a function of the preceding reward distribution, the GAC served to position utilities elicited in distributions with low and high means relative to adaptive or absolute utilities elicited from the full distribution.

The SAC represents the percent change in the CRs (the normalized areas under each curve) of successive utilities. It can be used to quantify differences in utilities within a single distribution, or, in this case, between the median utilities of different distributions. Importantly, the SAC allowed us to quantify utility adaptation on a normalized scale: if utility patterns were fully adapting (i. e. fixed shape regardless of the distribution), the SAC would gravitate to 0. On the other hand, the SAC would become negative if utilities became more convex (since the area under the utilities would become smaller), and more positive if utilities became more concave. The other coefficient, the GAC, compared the utility of the low- and high-distributions with the full reward distribution’s utility function (Fig. 2b, dashed lines). Using the full-distribution utility as the ‘default’ utility shape, the GAC measured how different narrow utilities were – ranging from no or 0% adaptation (i. e. narrow utilities were but segments of an absolute full-distribution utility) to 100% adaptation (the utilities had a fixed form that simply adapted to new distributions). We used DCM-derived utilities to calculate these adaptation coefficients.

Using the SAC to quantify how median utilities changed between distributions, we found that the differences between utilities of Monkey A amounted to SACs of 0.37 and 0.35 for the full- and high-distributions, respectively; 0.11 and -0.14 for Monkey B’s high- and full-distribution, and 0.04 for Monkey C’s low distribution. In utility terms, this meant that Monkey A’s utilities predicted behaviour that was 37% and 35% more risk averse in consecutive distributions. Monkey B also became more risk averse when going from the low distribution to the high distribution but became more risk seeking again once choosing in the full distribution. The direction of these changes seemed to reflect the ‘position’ of the tested distributions relative to the past distributions the animals had experienced. In line with this idea, Monkey C had no recent experience with the full-distribution when low-distribution utilities were estimated; the measured utilities were thus almost identical.

The GACs calculated for each animal were also very informative in positioning low- and high-distribution utilities relative to the full distribution ones (see dotted lines in Fig. 7). Monkey A, for example, had a GAC of 0.51 for the small distribution, and a GAC of 0.21 for the high distribution. The high GAC essentially meant that the low-distribution utility was halfway between being only a segment of a fixed full-distribution utility and being a fully rescaled versions of the full-distribution utility; the low GAC suggested that high-distribution utilities were much closer to being segments of a larger, absolute utility function. For Monkey B, low-distribution utilities matched a GAC of 1.14, i.e. The utilities of the low distribution had an almost identical shape to those in the full-distribution, and the high-distribution utilities had a GAC of 0.69, a bit more than halfway between no- and full-adaptation. Monkey C, corroborating earlier findings, had a GAC between low and full-distributions of 0.87 – they were, for all intents and purposes, identical.

Finally, going back to the original idea that preferences are shaped by one’s expectations, we looked at the shape of each DCM-utility relative to the task’s daily reward statistics. Though even the initial distribution’s utility inflections never truly followed the task’s mean reward (one-sample t-test; Monkey A: t(20)_low-distribution_ = 3.849, p = 0.001; Monkey B: t(23)_full-distribution_ = 2.534, p = 0.019; Monkey C: t(13)_high-distribution_ = 4.267, p = 1.103 x 10^−4^), the difference between mean rewards and inflections became markedly larger for Monkeys A and B when they were introduced to new reward distributions (Kruskal-Wallis test; Monkey A: H(2,58)= ‘40.052, p = 2.008 x 10^−9^; Monkey B: H(2,40)= ‘16.806, p = 2.242 x 10^−4^). Importantly, the differences were always skewed towards past distributions. As reward distributions changed, Monkey A and B’s references appeared to lag in fully adapting to the new distributions. Monkey C, on the other hand, saw no differences between its two reward distributions (H(1,54) = 0.021, p = 0.884) – presumably because of the 54-week gap between the two sets of measurements.

To better understand and quantify the lag in fully adapting to current reward, we built a simple reinforcement-learning model that predicted the reward distributions most likely to have shaped animals’ utilities (Sutton & Barto, 2018). Assuming the ‘normal’ form and a simple Rescorla–Wagner learning rule, the model then identified the distributions closest to the one captured by animal’s daily utility measures (that is, seeing utilities as the cumulative representation of the reward distribution the animals most expected). These distributions’ means and standard deviations (STD) were given by the following rule:

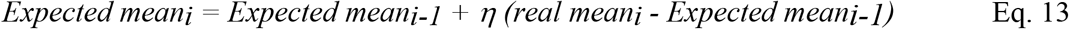

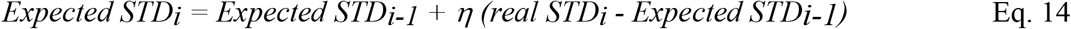

where each day’s ‘*expected*’ distribution relied on predictions from the previous day (_i-1_), as well as the learning rate (η) at which animals learn from the difference between these predictions (*expected*_i-1_) and reality (*real*_i_) – the prediction error. Importantly. the first *expected* parameters were assumed to be the statistics that the animal first observed, because of this as h would get closer to 1, it would indicate that predictions adapted instantly to new distributions; if h was closer to 0, it indicated preferences had relied only on early observations (i. e. the first distribution of rewards that the animal experience). The functions were fitted by minimizing the sums of square differences between the cumulative distribution function of these curves and the utility of the CEs that had been previously measured using the fractile method.

This simple reinforcement model offers insight as to the role that expectations played in shaping the animals’ preferences. Monkeys A, B, and C had learning rates of 0.62, 0.81, and 0.62, respectively; that is, their preferences adapted quickly to new reward distributions, but not fully. The recent past also played a role, albeit marginal, in shaping the relative value of rewards. Figure 9 illustrates both these ‘expected’ distributions as well as the ‘true’ distributions (as measured by the first derivative of the utility functions). Notice how the expected distributions spill over reward distribution changes only for the first couple of days. If preferences are built around expectations, then the utilities that best described these preferences point to these animals using mostly present but also past information to shape them.

**Figure 9.**
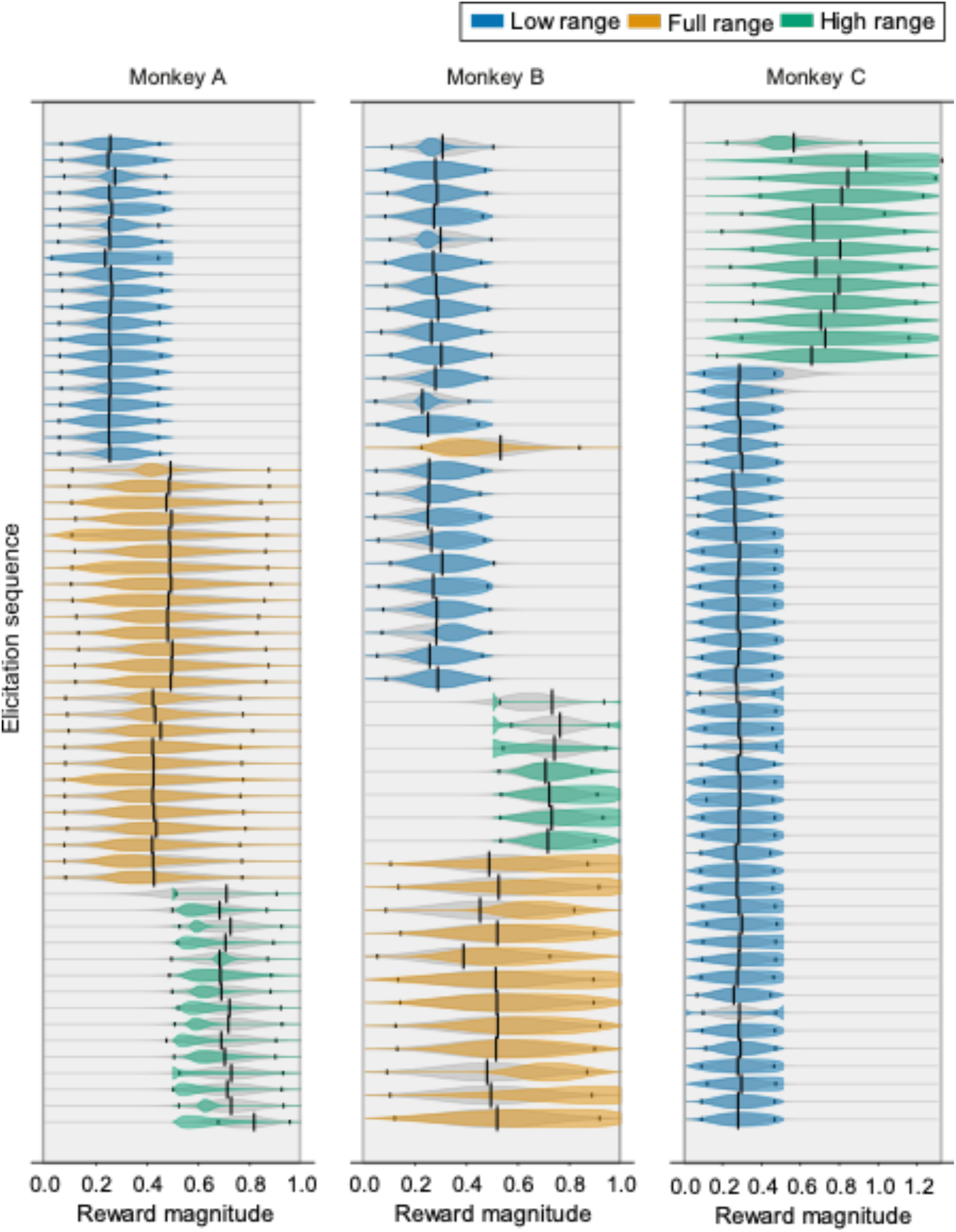
Daily inflections in utilities reflect recently experienced reward distributions. Each experimental session is represented by a set of horizontal black lines; the first derivatives of fitted utilities appear as the coloured ‘violin plots’ on the horizontal lines. Black vertical lines indicate the true mean of the rewards experienced by the animals on individual days – smaller black lines indicate the STD on these means. Grey ‘violin plots’ reflect the expected distributions of rewards that the animals ‘learned’ over past experimental sessions, based on reinforcement learning predictions (Eqs. 13; 14). They are the distributions that best fit utilities, as allowed by a Rescorla-Wagner learning rule. Of note, the grey normal distributions are not restricted by reward distributions in the way that utilities are.

### Discussion

The present study investigated the role of task-specific expectations in shaping the preferences of macaque monkeys. In line with human research on reference-dependent preferences (Arkes et al., 2008, 2010; Koszegi & Rabin, 2007), the animals’ risk preferences shifted following changes to the reward distribution they could expect from the task at hand. As the rewards that the task delivered got higher, the reward magnitude at which their risk-attitudes shifted also became higher. Modelling the utility functions that best captured the animals’ behaviour, we found that changes in their risk-preferences mimicked the changes predicted in models like Prospect Theory (Kahneman & Tversky, 1979): the points at which utility shifted from convex to concave closely followed what could be considered plausible expectations in the task.

Taking the position of S-shaped utilities as a proxy for the animal’s expectations, our findings suggest that the monkeys partially adapted their preferences to account for new reward distribution in a task. While they readily adapted to novel rewards, they did not readily ignore (or forget) reward information that was no longer relevant to the task. Rather than relying solely on the current instalment of the task to build their expectation, the monkeys appeared to also consider the distribution of past rewards – particularly the extremes in a distribution - in shaping their preferences (i. e. their utility curve). This led to partial, not full, adaptation.

Monkeys A and B, for example, reliably shifted their reference point when possible rewards went from lower to higher magnitudes. When looking at the utility function that best represented their preferences, the animals’ utilities appeared to scale instantly to represent the now broader realm of possible rewards. Conversely, when possible rewards were restricted to high magnitudes only (i.e. high-distribution), the animals did not adjust their preferences in a way that accounted for the unavailability of lower magnitudes – even after many days. Where they had previously been flexible in rescaling preferences, the animals’ preferences in the high distribution (where low rewards were never delivered) stubbornly reflected the higher-half of full-distribution utilities. And while the shift from low to high distribution seemed to induce partial, almost full adaptation – the shift from full to high distribution reflected a move along a fixed, absolute utility instead. The data from Monkey C, where different reward distributions were tested 54 weeks apart, corroborated this expectationbased interpretation by providing a window on the adaption of utilities after a year. While Monkeys A and B experienced every distribution in the span of just a couple of months, the effects of past high rewards on Monkey C would have been minimal. In that respect, it came as no surprise that Monkey C’s lower distribution utilities took the form of fully rescaled full-distribution ones. A similar effect was seen in previous estimations with Monkey A’s utilities (Genest et al., 2016).

The idea that preferences adapt to fit a given distribution is neither new nor unfounded (Brunswik, 1956; Gigerenzer et al., 1991; Glöckner et al., 2014; Weber & Johnson, 2008). Indeed, while prospect theory rests on reference-dependence, several newer models mimic RDU in that they claim that the values with which we imbue our options rely on the other options we have at our disposal (Hunter & Gershman, 2018; Loomes & Sugden, 2006; Parducci, 2012; Steward et al., 2003; Yaari, 2006). Likewise, it has long been known in psychology and neuroscience that distribution-adaptation is an inherent feature of the brain (Louie & De Martino, 2013). In sensory systems, for example, neuron’s maximize their efficiency by tuning their firing rates to match the distribution of sensory signals (Carandini & Heeger, 2012; Laughlin, 1981) – the same is thought to occur, to varying degrees, in the brain areas that encode value (Burke et al., 2016; Kobayashi et al., 2010; Louie et al., 2015; Padoa-Schioppa, 2009; Tobler et al., 2005; Tremblay & Schultz, 1999). Specifically, and supporting the idea of distribution-dependent utility, neurons in the primate prefrontal cortex have recently been recorded adapting their firing rate to different reward distributions in a way similar to our animals’ utility curves. In a study by Conen and Padoa-Schioppa (2019), rhesus macaques only partially rescaled the value of juice rewards relative to the other possibilities in a given block of choices. When recording from neurons in monkey orbitofrontal cortex, the researchers found that the neural code mimicked behavioural measurements in that it partially adapted to match the specific reward distributions of different blocks within the broader context of all past rewards. Crucially, two processes seemed to drive this adaptation: the first, a slow and adaptive learning process about the outcomes one can expect (e.g., reinforcement learning (Bavard et al., 2018; Rudebeck & Murray, 2014; Wilson et al., 2014), which involves the orbitofrontal cortex and its interaction with the dopaminergic system (for review, see Soltani & Izquierdo, 2019) and might explain the role of experience in shaping current preferences. The second process involves a rapid weighing of rewards relative the decision-maker’s present context (e.g., the canonical process of divisive normalization, whereby neurons tune their firing rates to match the distribution of available stimuli; Louie et al., 2013; Hiroshi Yamada, Louie, Tymula, & Glimcher, 2018; Zimmermann et al., 2018).

Partial adaptation is likely to underlie the brain’s ability to maximize ‘local’ decisions, all while placing these decisions in a much broader context (i.e. relative past experiences; Conen & Padoa-Schioppa, 2019; Fairhall, Lewen, Bialek, & De Ruyter van Steveninck, 2001; Rustichini, Conen, Cai, & Padoa-Schioppa, 2017). When comparing similarly-priced wines, for example, we manage to select our favourite from relatively narrow distributions (similar prices) while still placing our selection relative to a much broader price distribution (our past experiences with wines). It has recently been suggested that this ability to flexibly optimize ‘local’ decisions while keeping track of past outcomes underlies the formation of cause-and-effect relationships in our thinking (Bavard et al., 2018). If this is the case, then the changes observed in our animals’ utility functions point to the animals building complex expectations, or an internal model, about the rewards they could get in the task at hand.

Overall, and in line with the current view from neuroeconomics, this study showed that the preferences of macaque monkeys’ scale in a way that reflects both inherent properties (and indeed limitations) of the brain and the statistics of the task at hand. Put most poetically by the economists Herbert Simon, our animals’ decision appeared *“… shaped by scissors whose two blades are the structure of the task environments and the computational capabilities of the actor*” (Simon, 1990, p.7). Perhaps it is long time we consider this in the models used to study choice.

## Acknowledgements

Funding by Wellcome Trust (WT 095495, WT 204811), ERC Advanced Grant (293549).

